# Mabs, a suite of tools for gene-informed genome assembly

**DOI:** 10.1101/2022.12.19.521016

**Authors:** Mikhail I. Schelkunov

## Abstract

**Motivation:** Despite constantly improving genome sequencing methods, error-free eukaryotic genome assembly has not yet been achieved. Among other kinds of problems of eukaryotic genome assembly are so-called “haplotypic duplications”, which may manifest themselves as cases of alleles being mistakenly assembled as paralogues. Haplotypic duplications are dangerous because they create illusions of gene family expansions and, thus, may lead scientists to incorrect conclusions about genome evolution and functioning.

**Results:** Here, I present Mabs, a suite of tools that serve as parameter optimizers of the popular genome assemblers Hifiasm and Flye. By optimizing the parameters of Hifiasm and Flye, Mabs tries to create genome assemblies with the genes assembled as accurately as possible. Tests on 6 eukaryotic genomes showed that in 6 out of 6 cases, Mabs created assemblies with more accurately assembled genes than those generated by Hifiasm and Flye when they were run with default parameters. When assemblies of Mabs, Hifiasm and Flye were postprocessed by a popular tool for haplotypic duplication removal, Purge_dups, genes were better assembled by Mabs in 5 out of 6 cases.

**Availability and implementation:** Mabs has been written in Python and is available at https://github.com/shelkmike/Mabs

## 1 Introduction

In recent years, sequencing technologies have improved significantly. Reads of Oxford Nanopore Technologies have become longer and more accurate (Wang *et al.*, 2021), as have HiFi reads of PacBio (Pacific Biosciences, 2019, 2021). Despite this progress, genome assemblies still suffer from a number of problems, among the major of which are:

1. Fragmentation owing to long repeats with similar copies (Dida and Yi, 2021; Nurk *et al.*, 2022).
2. Contamination (Steinegger and Salzberg, 2020; Cornet and Baurain, 2022).
3. Haplotypic duplications (Guan *et al.*, 2020; Ko *et al.*, 2022).

The latter problem is a case where, during assembly of a diploid or polyploid genome, a genome assembler mistakes corresponding regions of two homologous chromosomes with regions that originated from segmental duplications. For example, two alleles of the same gene may be mistaken for paralogues. Haplotypic duplications are dangerous because they may lead to incorrect scientific conclusions about the gene content of a genome. When two alleles are assembled separately as paralogues, an illusion of a gene duplication event is created. In highly heterozygous genomes, haplotypic duplications are so frequent that they can result in such false duplicates for thousands of genes (Guan *et al.*, 2020; Ko *et al.*, 2022).

One way to address haplotypic duplications is to minimize them during the process of genome assembly. For example, authors of the genome assembler Hifiasm endowed it with a special algorithm that distinguishes corresponding regions of homologous chromosomes and segmental duplications (Cheng *et al.*, 2021). An alternative method is to try to remove haplotypic duplications after assembly. This method is implemented in specialized programs, such as Purge_dups (Guan *et al.*, 2020), Purge_haplotigs (Roach *et al.*, 2018) and HapSolo (Solares *et al.*, 2021). However, there are no methods or combinations of methods that are 100% effective in the removal of haplotypic duplications.

To address the problem of haplotypic duplications, I created a suite of tools called “Mabs” that I describe in this article. The main two components of Mabs are Mabs-hifiasm and Mabs-flye, which serve as parameter optimizers of the popular genome assemblers Hifiasm (Cheng *et al.*, 2021) and Flye (Kolmogorov *et al.*, 2019). Mabs-hifiasm is intended for assembly using PacBio HiFi (also known as PacBio CCS) reads, while Mabs-flye is intended for assembly using reads of more error-prone technologies, namely, Oxford Nanopore Technologies and PacBio CLR. By optimizing the parameters of Hifiasm or Flye, Mabs reduces the number of haplotypic duplications.

## 2 Materials and Methods

### 2.1 Overview of the algorithm

For a detailed description of the algorithm, see Supplementary Text S1.

Mabs tries to find values of parameters of a genome assembler that maximize the number of accurately assembled BUSCO genes. BUSCO is a program that is supplied with a number of taxon-specific datasets that contain orthogroups whose genes are present and single-copy in at least 90% of genomes from a given taxon (Simão *et al.*, 2015). Thus, it is expected that most BUSCO genes in any newly studied genome will also be present and single-copy. Mabs discriminates BUSCO genes into three categories:

1. Genes belonging to single-copy orthogroups.
2. Genes belonging to false multicopy orthogroups. These genes are haplotypic duplications.
3. Genes belonging to true multicopy orthogroups. These genes are paralogues.

The original algorithm of BUSCO does not differentiate between false multicopy and true multicopy orthogroups, combining all orthogroups into a single category “D” (short for “duplicated”). However, to properly evaluate the quality of a genome assembly, it is preferential to be able to distinguish false multicopy orthogroups from true multicopy orthogroups. To do this, Mabs uses the following procedure (see also Fig. 1A):

1. It calculates the median coverage in exons of BUSCO genes from single-copy orthogroups. I denote it Cov(S).
2. It calculates the median coverage in exons of BUSCO genes from multicopy orthogroups. The coverage is calculated separately for each BUSCO orthogroup.
3. If the median coverage in exons of genes of a multicopy BUSCO orthogroup is closer to Cov(S) than to (Cov(S)/2), then genes of this orthogroup are considered true multicopy genes. Otherwise, they are considered false multicopy genes. This method uses the fact that the coverage of alleles not collapsed during the assembly should be two times lower than the coverage of collapsed alleles.

**Fig. 1.**
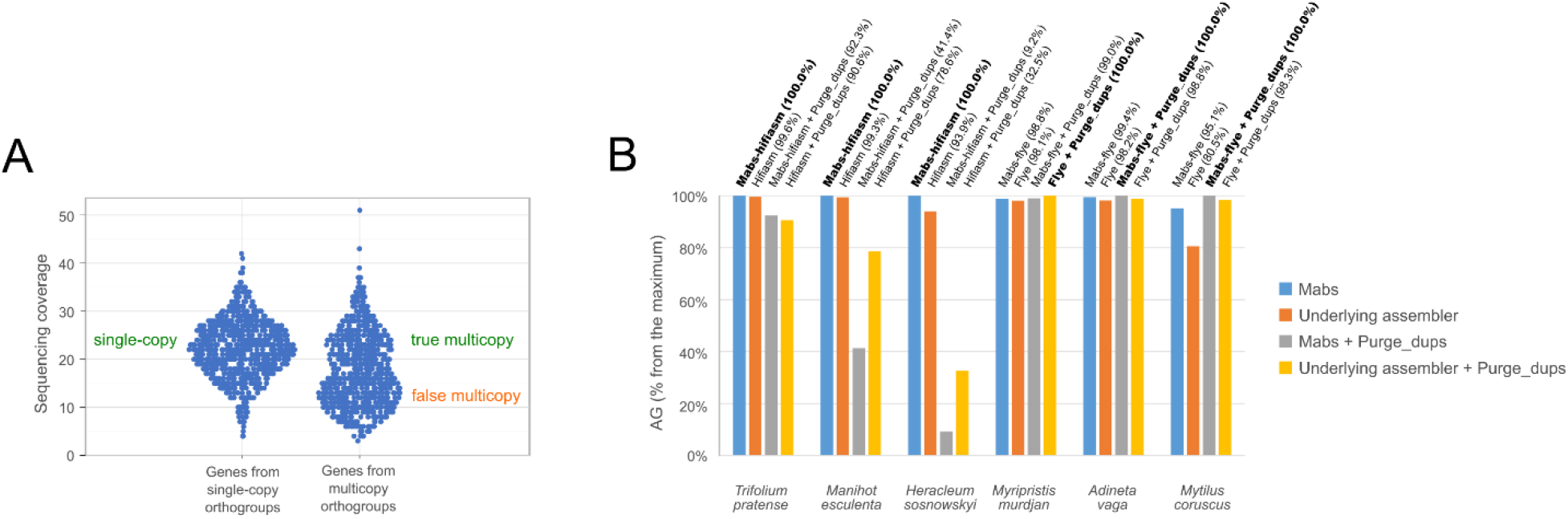
(**A**) Sinaplots (Sidiropoulos *et al.*, 2018) with coverage of BUSCO genes in the Hifiasm assembly of the genome of *Heracleum sosnowskyi*. Every point is a gene. For multicopy orthogroups, separation into true multicopy (high coverage) and false multicopy (low coverage) can be seen. (**B**) Values of AG for assemblies of the six genomes. AG is indicated percentage with respect to the maximum value achieved for this genome. The best assemblies are marked in bold.

Mabs uses a metric that I call AG (short for “Accurately assembled Genes”), which is calculated as the sum of the number of BUSCO genes in single-copy orthogroups and the number of BUSCO genes in true multicopy orthogroups. Basically, Mabs works as a parameter optimizer of a genome assembler, trying to find such values of parameters that maximize AG. An underlying assumption of Mabs is that optimization of the assembly quality of BUSCO genes also leads to optimization of the assembly quality of other protein-coding genes. Thus, by maximizing AG, Mabs tries to create a genome assembly where protein-coding genes are assembled as accurately as possible.

In principle, this strategy of “gene-informed parameter optimization” may be applied to any genome assembler that has some parameters that affect its algorithm. I have chosen Hifiasm and Flye because in multiple benchmarks, they were shown to be the best or among the best assemblers for accurate (PacBio HiFi) and error-prone (Oxford Nanopore Technologies and PacBio CLR) reads, respectively (Jung *et al.*, 2020; Murigneux *et al.*, 2020; Dida and Yi, 2021; Gavrielatos *et al.*, 2021; Guiglielmoni *et al.*, 2021; Schneider *et al.*, 2021; Wick and Holt, 2021; Rabanal *et al.*, 2022; Xie *et al.*, 2022; Zhang *et al.*, 2022). Since Hifiasm and Flye were favourably compared with other genome assemblers many times, in this work, I compare Mabs only with Hifiasm and Flye and do not make comparisons with other assemblers.

### 2.2 Genomes

To test Mabs, I performed a literature analysis and compiled a set of 5 species for which authors reported a high number of haplotypic duplications: *Trifolium pratense* (Bickhart *et al.*, 2022), *Manihot esculenta* (Qi *et al.*, 2022), *Myripristis murdjan* (Guan *et al.*, 2020; Rhie *et al.*, 2021), *Adineta vaga* (Guiglielmoni *et al.*, 2021), and *Mytilus coruscus* (Li *et al.*, 2020; Sun *et al.*, 2021). In addition, I included *Heracleum sosnowskyi*, which I studied personally and which also had many haplotypic duplications (to be published). For detailed information about the genomes and their sequencing reads, see Supplementary Tables S1 and S2. The genomes of *Trifolium pratense*, *Manihot esculenta* and *Heracleum sosnowskyi* were sequenced using PacBio HiFi technology and were thus used to compare Mabs-hifiasm and Hifiasm. The other three genomes were sequenced using error-prone technologies and were thus used to compare Mabs-flye and Flye.

## 3 Results

A detailed comparison of assemblies made by Mabs and its underlying assemblers Hifiasm and Flye is provided in Supplementary Table S3.

If AG was used as the metric of gene assembly quality, for all 6 genomes, Mabs created better assemblies than Hifiasm and Flye (Fig. 1B). A larger AG implies a higher quality of assembly of protein-coding genes. The larger amount of haplotypic duplications in assemblies made by Hifiasm and Flye can also be seen as larger values of BUSCO's “D” (the number of multicopy orthogroups), see Supplementary Table S3. In several of the most prominent cases, the larger amount of haplotypic duplications made by the underlying assemblers compared with Mabs can be seen through diagrams of gene coverage (see Supplementary Figs. S1 and S2). In terms of N50, assemblies made by Mabs were better than assemblies made by Hifiasm and Flye for 5 of 6 genomes.

When Purge_dups was applied after the assembly, assemblies made by Mabs were better than assemblies of Hifiasm and Flye for 5 of 6 genomes (Supplementary Table S3). Overall, Purge_dups was detrimental (it decreased AG) for all assemblies from HiFi reads but was beneficial for all assemblies from error-prone reads.

Despite the higher assembly quality, for all 6 genomes, Mabs was slower than the underlying assemblers, with the assembly time of Mabs being 1.6-3.5 times longer than those of the underlying assemblers. RAM consumption was approximately the same as that of the underlying assemblers (Supplementary Table S3).

## 4 Conclusions

In this article, I demonstrated the utility of the tool Mabs that optimizes the parameters of the genome assemblers Hifiasm or Flye. The idea of “gene-informed parameter optimization” used by Mabs may potentially be implemented with other genome, metagenome or transcriptome assemblers as well.

## Acknowledgements

The author is thankful to Maria D. Logacheva, Aleksey A. Penin, Artem S. Kasianov and Maksim S. Makarenko for their valuable comments.

The development of Mabs was funded by the budgetary subsidy with the code FFNU-2022-0037 to the Institute for Information Transmission Problems of the Russian Academy of Sciences. The genome assembly for *Heracleum sosnowskyi* was funded by research grant no. 21-74-20145 of the Russian Science Foundation.

