## Supplementary figures and tables for "Mabs, a suite of tools for gene-informed genome assembly"

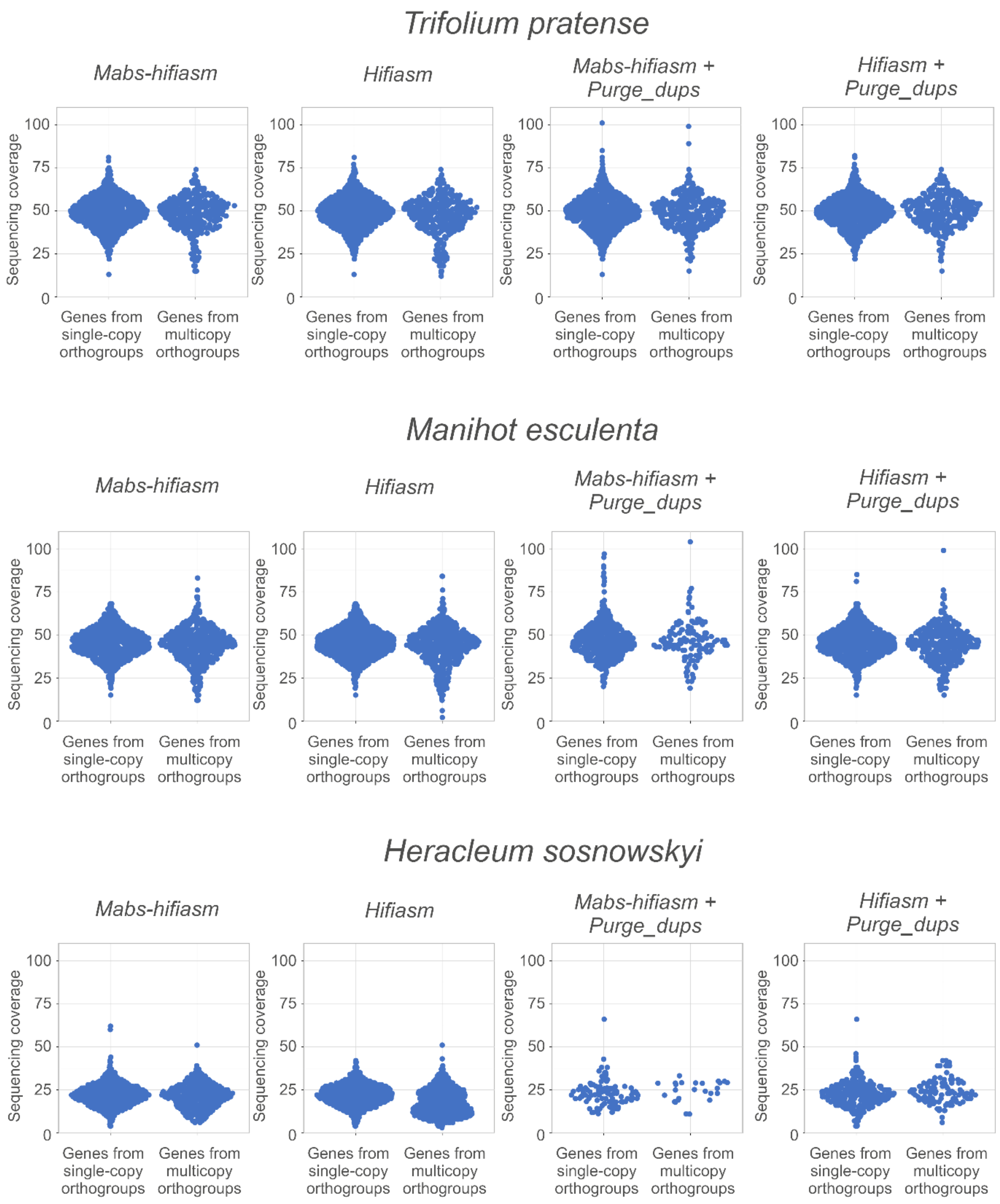


**Supplementary Figure S1. Sinaplots of BUSCO genes' coverage in assemblies of Mabs-hifiasm and Hifiasm**. Each dot is a BUSCO gene. More accurate removal of haplotypic duplications by Mabs-hifiasm in comparison with Hifiasm can be seen, for example, as the lower number of dots in the right bottom corner in the first two diagrams for *Heracleum sosnowskyi*.

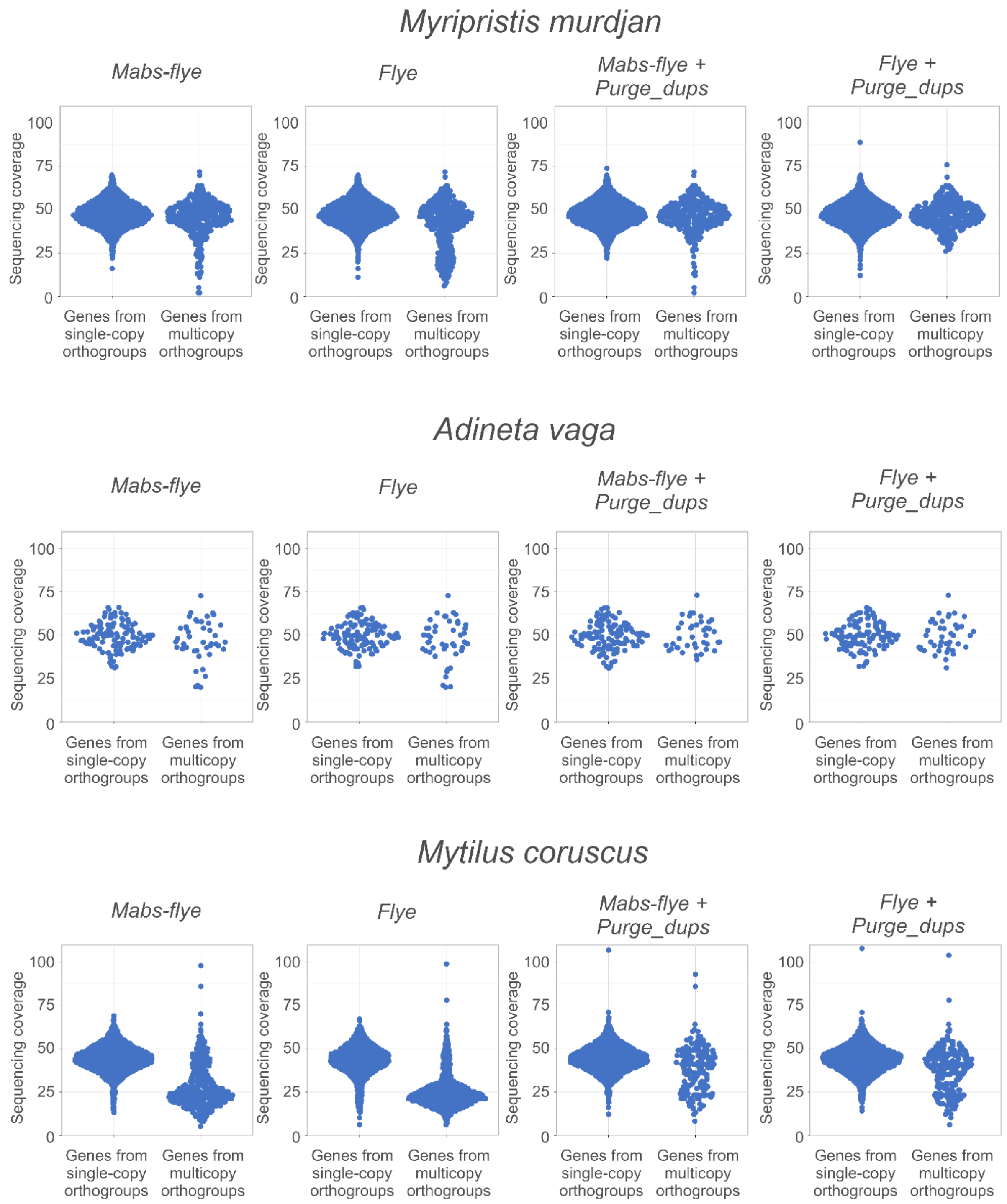


**Supplementary Figure S2. Sinaplots of BUSCO genes' coverage in assemblies of Mabs-flye and Flye.** Each dot is a BUSCO gene. More accurate removal of haplotypic duplications by Mabs-flye in comparison with Flye can be seen, for example, as the lower number of dots in the right bottom corner in the first two diagrams for *Myripristis murdjan*.

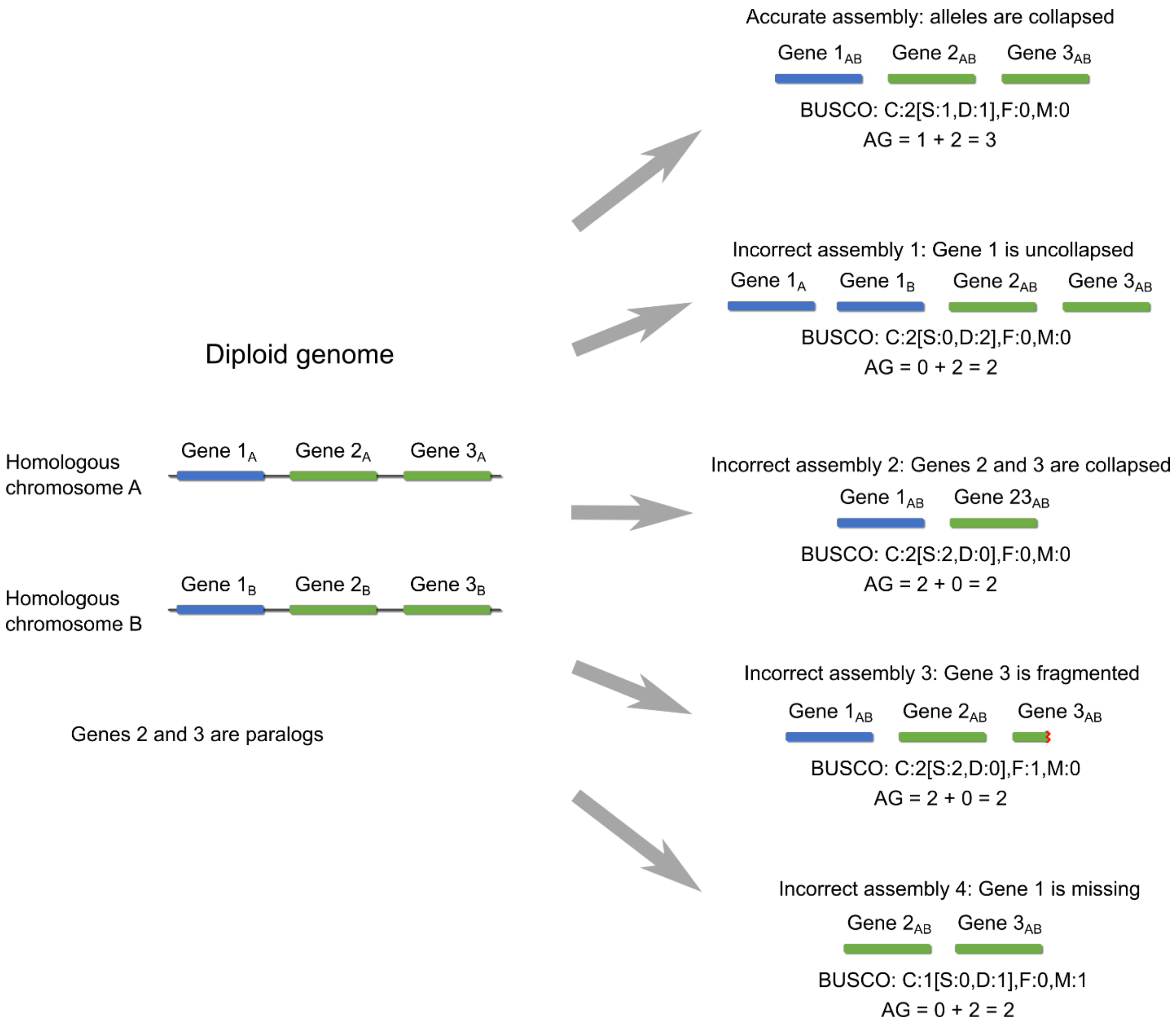


**Supplementary Figure S3. Some of the ways a gene assembly may fail.** In this simplified scheme, three BUSCO genes are depicted, two of them are paralogs. BUSCO completeness ("C") is equal in the first four assemblies; however, only one of these assemblies is correct. At the same time, the largest AG clearly defines the best assembly. Other variants of gene misassembly are also possible, but not shown.

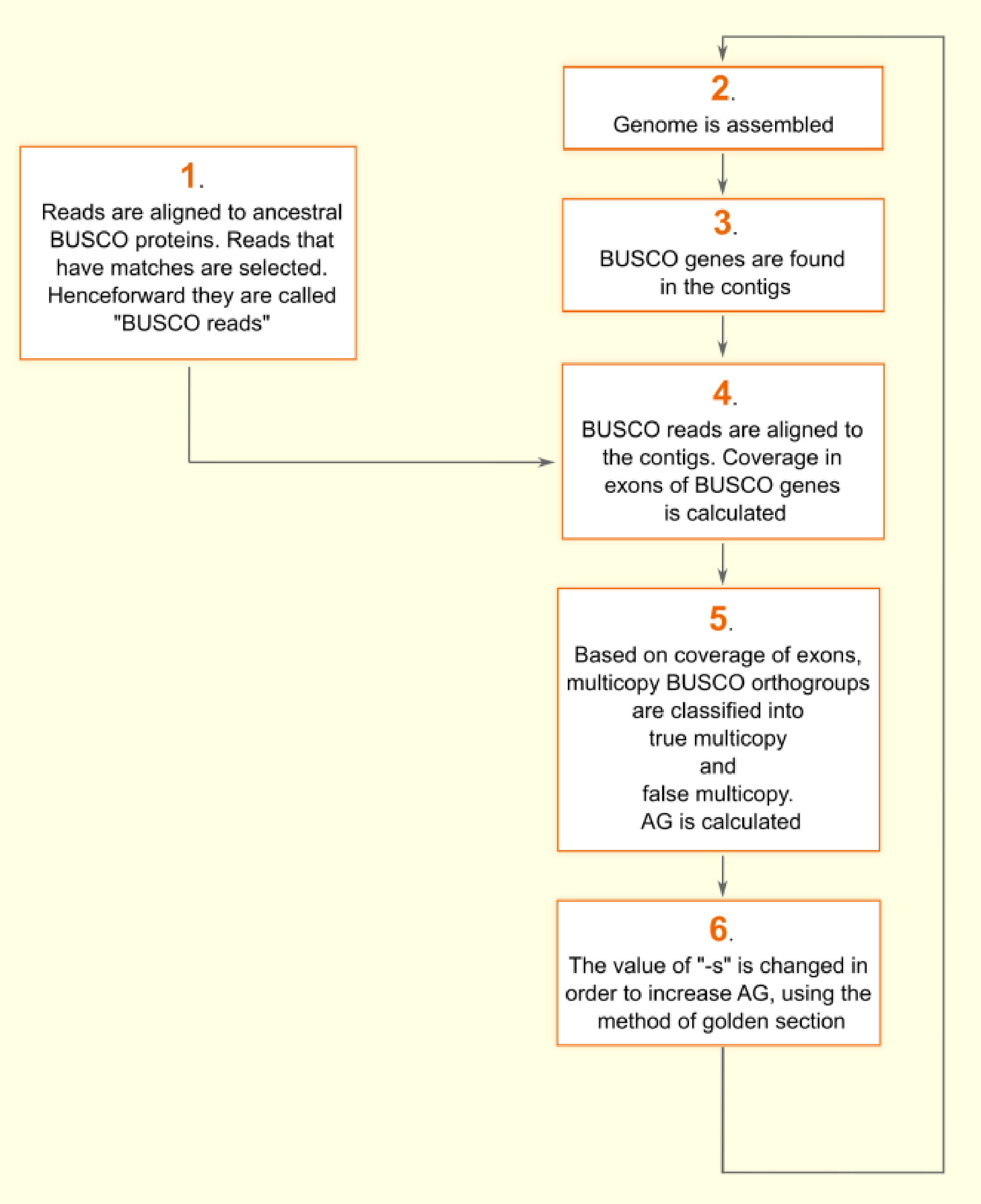


**Supplementary Figure S4. The flowchart of Mabs-hifiasm**

| Species | Genome size, estimated without sequencing (Mbp) | Genome size, estimated as the size of the assembly with the largest N50 among published assemblies (Mbp) |
|---|---|---|
| *Trifolium pratense* | 636 (Grime and Mowforth, 1982), 474 (Arumuganathan and Earle, 1991), 418 (Vižintin *et al.*, 2006), 557 (Zonneveld, 2019), | 414 (Bickhart *et al.*, 2022) |
| *Manihot esculenta* | 817 (Awoleye *et al.*, 1994) | 706 (Qi *et al.*, 2022) |
| *Heracleum sosnowskyi* | 1751 (Zonneveld, 2019) | 1629 (preprint to be uploaded to BioRxiv in 2023) |
| *Myripristis murdjan* | - | 835 (Vertebrate Genomes Project) |
| *Adineta vaga* | 362 (Welch and Meselson, 2003) | 101 (Simion *et al.*, 2021) |
| *Mytilus coruscus* | 1858 (Ieyama *et al.*, 1994) | 1567 (Yang *et al.*, 2021) |

**Supplementary Table S1. Information about genomes used in the article.**

| Species | Technology | Sequence Read Archive identifier | N50 of reads (bp) | Approximate genome coverage [a] |
|---|---|---|---|---|
| *Trifolium pratense* | PacBio HiFi | SRR15433789 | 20,082 | 50 |
| *Manihot esculenta* | PacBio HiFi | ERR5485301 | 20,363 | 42 |
| *Heracleum sosnowskyi* | PacBio HiFi | SRR23251371, SRR23251372 | 14,679 | 22 |
| | Illumina Hi-C, paired-end | SRR23251383, SRR23251384 | 76 | 34 |
| *Myripristis murdjan* | PacBio CLR | ERR3449630, ERR3449634, ERR3449635, ERR3453872, ERR3453873, ERR3453874, ERR3453875, ERR3453876 | 24,966 | 50 |
| | Illumina shotgun, paired-end [b] | ERR3655549 | 151 | 155 |
| *Adineta vaga* | Oxford Nanopore | SRR13348928 | 38,562 | 50 |
| | Illumina shotgun, paired-end [b] | SRR13348929 | 251 | 331 |
| *Mytilus coruscus* | Oxford Nanopore | ERR3415816 | 24,641 | 50 |
| | Illumina shotgun, paired-end [b] | ERR3431204 | 150 | 53 |

**Supplementary Table S2. Information about reads used in the article to assemble the genomes.** Reads with coverage 50 were downsampled from larger amounts of reads by Filtlong to accelerate assembly.

[a] the approximate genome coverage was calculated based on the following genome sizes: *Trifolium pratense* 450 Mbp, *Manihot esculenta* 750 Mbp, *Heracleum sosnowskyi* 1700 Mbp, *Myripristis murdjan* 850 Mbp, *Adineta vaga* 100 Mbp, *Mytilis coruscus* 1600 Mbp.

[b] used only for polishing, after the assembly.

| Species | Method of assembly | BUSCO results | N50 (bp) | Sum of contigs' lengths (bp) | AG | Assembly time[a] | Peak RAM usage[a] |
|---|---|---|---|---|---|---|---|
| *Trifolium pratense* | Mabs-hifiasm | C:98.0%[S:92.9%,D:5.1%], F:1.3%,M:0.7% | 20,490,459 | 460,521,547 | 1385 | 9h 50m | 62 GB |
| | Hifiasm | C:98.1%[S:92.6%,D:5.5%], F:1.3%,M:0.6% | 18,371,892 | 472,503,778 | 1380 | 2h 55m | 54 GB |
| | Mabs-hifiasm + purge_dups | C:90.7%[S:86.3%,D:4.4%], F:1.1%,M:8.2% | 21,823,263 | 332,714,522 | 1279 | 9h 50m + 1h 41m | 62 GB |
| | Hifiasm + purge_dups | C:90.6%[S:86.2%,D:4.4%], F:1.4%,M:8.0% | 21,823,263 | 317,658,493 | 1255 | 2h 55m + 1h 45m | 54 GB |
| *Manihot esculenta* | Mabs-hifiasm | C:98.5%[S:91.0%,D:7.5%], F:0.7%,M:0.8% | 34,338,427 | 747,732,950 | 1590 | 10h 51m | 102 GB |
| | Hifiasm | C:98.5%[S:90.0%,D:8.5%], F:0.7%,M:0.8% | 29,220,819 | 774,580,850 | 1579 | 4h 18m | 94 GB |
| | Mabs-hifiasm + purge_dups | C:46.0%[S:42.8%,D:3.2%], F:1.0%,M:53.0% | 33,797,513 | 208,140,594 | 658 | 10h 51m + 5h 7m | 102 GB |
| | Hifiasm + purge_dups | C:80.5%[S:74.8%,D:5.7%], F:0.8%,M:18.7% | 31,723,266 | 455,473,424 | 1250 | 4h 18m + 5h 51m | 94 GB |
| *Heracleum sosnowskyi* | Mabs-hifiasm | C:98.0%[S:89.6%,D:8.4%], F:0.4%,M:1.6% | 22,232,970 | 1,631,972,638 | 1392 | 13h 45m | 57 GB |
| | Hifiasm | C:98.3%[S:80.9%,D:17.4%], F:0.5%,M:1.2% | 13,468,048 | 1,814,805,347 | 1307 | 4h 8m | 74 GB |
| | Mabs-hifiasm + purge_dups | C:11.4%[S:10.8%,D:0.6%], F:0.9%,M:87.7% | 55,489,431 | 178,451,278 | 128 | 13h 45m + 12h 33m | 57 GB |
| | Hifiasm + purge_dups | C:35.7%[S:34.0%,D:1.7%], F:0.8%,M:63.5% | 12,038,317 | 460,607,932 | 453 | 4h 8m + 16h 32m | 74 GB |
| *Myripristis murdjan* | Mabs-flye | C:97.2%[S:95.7%,D:1.5%], F:1.1%,M:1.7% | 1,738,191 | 849,324,108 | 2454 | 53h 14m | 18 GB |
| | Flye | C:97.9%[S:94.6%,D:3.3%], F:1.0%,M:1.1% | 1,831,223 | 913,966,709 | 2437 | 20h 15m | 25 GB |
| | Mabs-flye + purge_dups | C:97.2%[S:96.1%,D:1.1%], F:1.1%,M:1.7% | 1,883,475 | 812,749,847 | 2457 | 53h 14m + 24m | 18 GB |
| | Flye + purge_dups | C:97.8%[S:96.7%,D:1.1%], F:1.0%,M:1.2% | 2,177,694 | 834,985,181 | 2483 | 20h 15m + 20m | 25 GB |
| *Adineta vaga* | Mabs-flye | C:66.3%[S:57.0%,D:9.3%], F:10.4%,M:23.3% | 2,333,984 | 112,876,648 | 163 | 9h 50m | 32 GB |
| | Flye | C:67.0%[S:57.5%,D:9.5%], F:9.9%,M:23.1% | 1,636,027 | 116,202,497 | 161 | 2h 43m | 36 GB |
| | Mabs-flye + purge_dups | C:66.3%[S:57.3%,D:9.0%], F:10.4%,M:23.3% | 2,698,890 | 106,057,374 | 164 | 9h 50m + 6m | 32 GB |
| | Flye + purge_dups | C:66.6%[S:57.3%,D:9.3%], F:9.9%,M:23.5% | 2,339,515 | 105,961,627 | 162 | 2h 43m + 5m | 36 GB |
| *Mytilus coruscus* | Mabs-flye | C:85.0%[S:78.4%,D:6.6%], F:4.2%,M:10.8% | 308,470 | 2,165,590,554 | 2040 | 114h 46m | 210 GB |
| | Flye | C:83.7%[S:66.6%,D:17.1%], F:4.4%,M:11.9% | 280,259 | 2,333,181,150 | 1726 | 70h 10m | 216 GB |
| | Mabs-flye + purge_dups | C:84.3%[S:82.0%,D:2.3%], F:4.2%,M:11.5% | 343,029 | 1,951,757,051 | 2144 | 114h 46m + 1h 55m | 210 GB |
| | Flye + purge_dups | C:82.4%[S:79.8%,D:2.6%], F:4.5%,M:13.1% | 320,773 | 1,962,315,961 | 2107 | 70h 10m + 1h 49m | 216 GB |

**Supplementary Table S3. Statistics of genome assemblies.**

[a] Genomes were assembled using 50 threads of Intel Xeon E7-4830 CPUs.

| **Category** | **One-letter abbreviation** | **Meaning** |
|---|---|---|
| Single-copy | S | Orthogroups that have a single completely assembled gene in the studied genome. A gene is considered completely assembled if its protein passed two criteria:<br>a) A criterion for sequence similarity to reference BUSCO proteins<br>b) A criterion for minimum length. |
| Duplicated | D | Orthogroups that have more than one completely assembled gene in the studied genome. Criteria for completeness are the same as described for "S". |
| Fragmented | F | Orthogroups that contain only genes that do not pass the criterion "b)" but pass the criterion "a)". Presence of such genes may be indicative of misassemblies that have led to gene fragmentation. Alternatively, such genes may indeed be shorter than reference genes. |
| Missing | M | Orthogroups for which no genes passing the criterion "a)" were found. |
| Complete | C | A compound category that is composed of orthogroups from category "S" and orthogroups from category "D" together. |

**Supplementary Table S4. Orthogroup categories used by BUSCO**

| Species | BUSCO dataset | Total number of orthogroups in the BUSCO dataset | Number of orthogroups used during the assembly | Number of orthogroups used to test final assembly quality |
|---|---|---|---|---|
| *Trifolium pratense* | eudicots_odb10 | 2326 | 1000 | 1326 |
| *Manihot esculenta* | eudicots_odb10 | 2326 | 1000 | 1326 |
| *Heracleum sosnowskyi* | eudicots_odb10 | 2326 | 1000 | 1326 |
| *Myripristis murdjan* | actinopterygii_odb10 | 3640 | 1000 | 2640 |
| *Adineta vaga* | metazoa_odb10 | 954 | 500 | 454 |
| *Mytilus coruscus* | mollusca_odb10 | 5295 | 1000 | 4295 |

**Supplementary Table S5. The split-up of BUSCO datasets into parts for assembly and testing.**
