## Supplementary text for "Mabs, a suite of tools for gene-informed genome assembly"

### Supplementary Text S1 (Supplementary Methods)

#### 1 How can the quality of a genome assembly be evaluated?

##### 1.1 General considerations

When optimizing the parameters of a genome assembler, it is important to reasonably select a metric of assembly quality that will be maximized during optimization. Many methods for evaluating the quality of a genome assembly exist, the most popular of which are probably calculating the N50 value and performing BUSCO analyses.

### 1.2 N50

N50 is calculated as the length of the largest contig (or scaffold) such that it and all contigs (or scaffolds) longer than it constitute at least half of the sum of the lengths of all contigs (or scaffolds). Basically, N50 is a metric of contig length. The downside of N50 is that genome assemblers sometimes make improper junctions, joining sequences when they should not be joined, thus inflating N50. A parameter optimizer that maximizes N50 will favour such improper junctions.

##### 1.3 BUSCO results

BUSCO is a program that is provided with many taxon-specific datasets (Simão *et al.*, 2015). Each of these datasets contains information about orthogroups (I refer to them as "BUSCO orthogroups") that have only one gene (I refer to them as "BUSCO genes") in genomes of at least 90% of species from a reference set of species of this taxon. An assumption on which a BUSCO analysis of the quality of a newly studied genome is based is that most BUSCO orthogroups will likely have a single gene. Thus, the number of BUSCO genes found in a genome may serve as a metric of assembly quality.

For example, the BUSCO dataset for land plants (embryophyta_odb10.2020-09-10) consists of 1614 orthogroups and was made based on a reference set of genomes of 50 species from this taxon.

BUSCO classifies orthogroups into 5 categories (see Supplementary Table S4).

Below, I use a designation of N("S") for the number of orthogroups in category "S", and designations for the other four categories follow the same naming structure. The number of orthogroups in each of the five categories can serve as a separate metric of genome assembly accuracy. Of these five metrics, the metric most often used for genome assembly assessment is probably N("C").

However, the existence of haplotypic duplications does not decrease N("C"), since category "D" is part of category "C". When dealing with the problem of haplotypic duplications, a better method is to maximize N("S") rather than N("C"), since haplotypic duplications move orthogroups from "S" to "D", thus decreasing N("S").

The early version of Mabs was designed to optimize the parameters of genome assemblers to maximize N("S"). However, I later realized that maximization of N("S") has a disadvantage because it favours assemblies where paralogues are merged. Indeed, if a genome assembler improperly merges paralogues into a single gene, this will lead to an increase in N("S"), while N("D") decreases and N("C") does not change. To address this problem, I created a novel metric that I call AG, which is short for "Accurately assembled Genes".

### 1.4 AG

Hereafter, "multicopy orthogroups" refers to what authors of BUSCO called "duplicated orthogroups". In my opinion, "multicopy orthogroups" is a better term since orthogroups in the BUSCO category "D" sometimes contain more than two genes.

The idea behind AG is that multicopy orthogroups (orthogroups from the BUSCO category "D") may be classified into true multicopy (I designate them "TM") and false multicopy (I designate them "FM") based on their coverage. Genome assemblers are usually made in such a way that they collapse two alleles into a single sequence during the process of genome assembly. If an allele is uncollapsed (i.e., a haplotypic duplication has occurred), then the read coverage of genes of this multicopy orthogroup will be twice as low as expected. This allows differentiating true multicopy (i.e., composed of paralogues) and false multicopy (i.e., composed of uncollapsed alleles) orthogroups. AG is calculated as a sum of the following two values:

1. The number of genes in single-copy ("S") orthogroups.
2. The number of genes in true multicopy ("TM") orthogroups.

Note that AG is not a sum of the numbers of orthogroups but a sum of the numbers of genes in them. This is because if only one gene is assembled in an orthogroup that has two paralogues, the number of orthogroups will not change (one orthogroup moves from "TM" to "S"), but the number of accurately assembled genes decreases by 1 (the number of "TM" genes decreases by 2 and the number of "S" genes increases by 1). Thus, basing AG on the number of genes in correctly assembled orthogroups is better than basing AG on the number of correctly assembled orthogroups itself.

For a comparison of AG and BUSCO statistics, see Supplementary Figure S3.

AG is a single value that, in my opinion, may serve as a very informative measure of how well a genome is assembled. In addition to Mabs-hifiasm and Mabs-flye, the Mabs suite of tools includes a third tool called "calculate_AG" that allows a user to calculate AG for any genome assembly. This tool may be useful to compare several genome assemblies made by different genome assemblers to decide which one is most accurate.

One disadvantage of AG is that it is poorly suited to compare two *nearly perfect* genome assemblies. For example, the recent telomere-to-telomere human genome assembly was made for the genome of a hydatidiform mole, which has an advantage for performing genome assembly in that it is nearly 100% homozygous (Nurk *et al.*, 2022). With HiFi reads or ultralong Oxford Nanopore Technology reads, it is possible to obtain genome assemblies that are accurate to the level of all protein-coding genes being assembled perfectly. The main problem with such assemblies is the difficulty in assembling tandem repeats with long monomers, such as centromeres and rDNA clusters (Nurk *et al.*, 2022; Rabanal *et al.*, 2022; Mc Cartney *et al.*, 2022). Thus, any assembly quality metrics that are based on how well protein-coding genes are assembled, be it AG or any of the 5 metrics of BUSCO, will usually be useless for comparing two nearly perfect assemblies.

##### 1.5 How AG is calculated

Given a set of reads and a genome assembly, the AG calculation procedure is as follows:

1. BUSCO genes in a genome are predicted using a method that is, basically, a simplified version of the method used by BUSCO. I intentionally simplified the technique of BUSCO to increase the speed of prediction at the cost of slightly decreased accuracy. The prediction is performed as follows:

   a. Potential BUSCO genes are predicted in the assembly by MetaEuk (Levy Karin *et al.*, 2020) using "ancestral" BUSCO proteins as a reference. The ancestral BUSCO proteins are reconstructed proteins of the last common ancestor of the species used to form the BUSCO dataset (for example, the last common ancestor of the 50 plant species of the dataset for land plants mentioned above). Sequences of the ancestral proteins are provided with each BUSCO dataset. The use of ancestral proteins as a reference to search for genes in modern species is beneficial because they are approximately equidistant to all modern species if the mutation accumulation rate did not differ greatly among lineages during evolution. In contrast, proteins of some modern genomes may be less suitable as a reference, since the phylogenetic distance to the genome under analysis may be larger and, thus, sequence similarity may be lower, making protein-to-genome alignment more difficult.
   b. Proteins of potential BUSCO genes predicted by MetaEuk are compared with profile Markov models of reference proteins from BUSCO orthogroups. This comparison is performed by the program "hmmsearch" from the HMMER suite of programs (Mistry *et al.*, 2013).
   c. To discriminate genes of BUSCO orthogroups from distant homologues, Mabs uses the same two criteria with the same threshold values as BUSCO: a) the criterion for sequence similarity to reference BUSCO proteins based on bit scores calculated by HMMER as described above and b) the criterion for minimum length.
2. Long reads are aligned to the genome by Minimap 2 (Li, 2018).
3. Sequencing coverage in exons of all identified BUSCO genes is calculated. Mabs does not calculate coverage in introns because introns may contain transposable elements. Copies of transposable elements may be assembled incorrectly in other regions of the genome, which may lead to distorted read coverage in introns. Distorted read coverage, in turn, may decrease the accuracy of classification of multicopy orthogroups into true multicopy and false multicopy.
4. The median coverage is calculated for all single-copy orthogroups. I denote it as Cov(S).
5. For each multicopy BUSCO orthogroup, an average value between median exonic coverages of all its genes is calculated. Since true multicopy orthogroups are likely to have coverage approximately equal to Cov(S) and false multicopy orthogroups (originating from haplotypic duplications) are likely to have coverage approximately equal to Cov(S)/2, Mabs uses a threshold of (3/4) * Cov(S) to discriminate true multicopy orthogroups from false multicopy orthogroups.
   Actually, the coverage distribution of genes with average coverage Cov(S)/2 may be narrower than the coverage distribution of genes with average coverage Cov(S) if the distribution behaves similarly to the Poisson distribution, where variance increases with increasing average. Hence, the threshold should probably be somewhat lower than (3/4) * Cov(S). The threshold was set to (3/4) * Cov(S) for simplicity.
6. AG is calculated as the sum of the number of genes in single-copy orthogroups and the number of genes in true multicopy orthogroups.

Some BUSCO datasets are composed of a very large number of orthogroups. For example, the dataset for primates (primates_odb10.2021-02-19) contains 13,780 orthogroups. Searching for genes of all these orthogroups in an assembly is time-consuming. At the same time, to estimate the quality of a genome assembly, a smaller number of orthogroups is probably sufficient. Hence, to save time, for any dataset that contains more than 1000 orthogroups, Mabs by default uses only 1000 orthogroups with the most conserved sequences. Orthogroups with the most conserved sequences are determined as orthogroups with the least mean positional relative entropy, as calculated by the program "hmmstat" from the HMMER suite of programs. The use of orthogroups with conserved sequences is preferential for genome assembly quality evaluation because it decreases the chance of genes not being identified because of too diverged sequences. Mabs has an option "--number_of_busco_orthogroups" that allows a user to set the number of BUSCO orthogroups to a value other than 1000.

### 2 Which parameters to optimize?

#### 2.1 General considerations

When making a parameter optimizer for a program, it is important to choose which parameters will be optimized.

From one perspective, optimizing too many parameters at the same time requires exploration of a multidimensional space of parameters, which demands considerable time. Exploration of a multidimensional space of parameters of a genome assembler is especially time-consuming because testing a single point in the space (i.e., performing one genome assembly) may take hours or days for a eukaryotic genome; see the assembly time for Hifiasm and Flye in Supplementary Table S3. For especially large genomes, assembly with a relatively slow genome assembler may take months (Neale *et al.*, 2022).

From another perspective, the more parameters are optimized, the greater the possible improvement in genome assembly.

Genome assemblers sometimes have dozens of parameters that affect their algorithm. The most prominent example that I have seen is Shasta (Shafin *et al.*, 2020), with Shasta 0.10.0 having 116 parameters that may affect the produced assembly.

#### 2.2 Hifiasm

For Hifiasm, the choice of a parameter for optimization is straightforward. It is the parameter "-s" that regulates the work of a special algorithm of Hifiasm made specifically to address haplotypic duplications. "-s" can have values in the range of 0 to 1. The default "-s" value in Hifiasm is 0.55, except when performing trio binning (usage of reads of both parents of the studied organism during assembly), where haplotypic duplication removal is not used. The algorithm behind "-s" is described in (Cheng *et al.*, 2021), but speaking simply, the closer the value of "-s" is to 0, the more aggressive Hifiasm is in the removal of similar sequences from the assembly.

The sole parameter of Hifiasm that Mabs-hifiasm optimizes is "-s".

#### 2.3 Flye

Optimization of Flye to reduce the number of haplotypic duplications is not as straightforward as the optimization of Hifiasm, since Flye has no parameters dedicated specifically for removal of haplotypic duplications, except for the parameter "--no-alt-contigs", which is logical ("true" or "false"). My tests (data not provided) indicate that "--no-alt-contigs" is probably always beneficial for removal of haplotypic duplications, thus Mabs-flye always runs Flye with this option. Based on my understanding of the algorithm of Flye, I chose two parameters for optimization:

1. "assemble_ovlp_divergence". When used in combination with "assemble_divergence_relative=0", as in Mabs-flye, the parameter "assemble_ovlp_divergence" regulates how dissimilar sequencing reads are allowed to be during disjointig construction. For a description of the algorithm and the term "disjointig" see (Kolmogorov *et al.*, 2019), but basically, higher values of "assemble_ovlp_divergence" may lead to more aggressive removal of similar sequences from the assembly.
2. "repeat_graph_ovlp_divergence". This parameter regulates how dissimilar sequences from disjointigs are allowed to be during merging of disjointigs into a repeat graph. As with "assemble_ovlp_divergence", larger values of this parameter may lead to more aggressive removal of similar sequences from the assembly.

Default values of these parameters in Flye differ depending on the sequencing technology used to produce the reads being assembled.

To accelerate the assembly, Mabs-hifiasm assumes these two parameters to be equal, referring to them as a single parameter "max_divergence". Thus, the parameter optimization is performed by Mabs-flye in a unidimensional space, just as in the case with Mabs-hifiasm.

### 3 The workflow of Mabs-hifiasm and Mabs-flye

#### 3.1 General considerations

The workflow of Mabs-hifiasm is similar to the workflow of Mabs-flye. In Section 3.2 I will describe the workflow of Mabs-hifiasm, and then, in Section 3.3, I will pinpoint the differences between Mabs-flye and Mabs-hifiasm.

#### 3.2 Mabs-hifiasm

The basic scheme of Mabs-hifiasm is provided in Supplementary Figure S4. Speaking simply, Mabs-hifiasm tries to find the value of the "-s" parameter of Hifiasm that provides as large an AG value as possible. The maximization is performed using the method of the golden section (Kiefer, 1953). The parameter "-s" can range from 0 to 1. In the golden section method, the first two values to be examined are middle points, $(\frac{\sqrt{5}-1}{\sqrt{5}+1})$ and $(1-\frac{\sqrt{5}-1}{\sqrt{5}+1})$, while the next values are determined based on the AG values.

The basic steps in the workflow of Mabs-hifiasm are as follows:

1. Reads are aligned to ancestral BUSCO proteins by DIAMOND (Buchfink *et al.*, 2015). The purpose is to select reads that belong to BUSCO genes. Hereafter I refer to them as "BUSCO reads". They will be used in step 4 to calculate the coverage of exons of BUSCO genes. Of course, it is possible to use all reads for the calculation of coverage, but using only BUSCO reads saves time since they constitute only a portion of all reads. For large eukaryotic genomes, where most regions are intergenic or intronic, the time saved by aligning only BUSCO reads becomes especially prominent.
   Since errors in long reads are often indels (Wenger *et al.*, 2019; Dohm *et al.*, 2020; Delahaye and Nicolas, 2021; Baid *et al.*, 2022) and, thus, may lead to frameshifts, DIAMOND is run with the option "--frameshift", which allows for frameshifts in the alignment.

2. A genome is assembled by Hifiasm using the current value of "-s".

3-5. AG is calculated as described in the section "**1.5 How AG is calculated**". This step is represented by three boxes (from "BUSCO genes..." to "Based on coverage of exons...") in Supplementary Figure S4.

6. Based on the AG value, the next value of "-s" is selected using the golden section method.

Ten points (including the two starting middle points) are examined by Mabs-hifiasm during the golden section optimization. Including more points may provide more accuracy at the cost of increasing assembly time. My tests show that 10 points is more than sufficient for determining the value of "-s" that provides the maximum or nearly maximum AG value.

##### 3.3 Mabs-flye

Mabs-flye uses basically the same workflow as Mabs-hifiasm, with the following differences:

1. Instead of Hifiasm, Mabs-flye uses Flye as the genome assembler. The need for two separate tools (Mabs-hifiasm and Mabs-flye) appeared because the algorithm of Hifiasm is intended foremost for very accurate (PacBio HiFi) reads, while the algorithm of Flye is intended mainly for considerably less accurate (PacBio CLR or Oxford Nanopore) reads. As of 2023, both PacBio HiFi and Oxford Nanopore technologies are widely used; thus, Mabs is split into Mabs-hifiasm and Mabs-flye. Although Flye has a dedicated option that allows it to assemble PacBio HiFi reads, "--pacbio-hifi", my tests on several genomes (data not provided) suggest that Hifiasm usually assembles genomes from HiFi reads better than Flye does.
   Taking into account the currently increasing accuracy of Oxford Nanopore reads (Wang *et al.*, 2021; Oxford Nanopore Technologies, 2022), it is possible that in the future, Hifiasm will also be suitable for Oxford Nanopore reads, thus reducing the necessity for Flye and, consequently, for Mabs-flye.
2. While Mabs-hifiasm optimizes the parameter "-s" of Hifiasm, Mabs-flye optimizes the parameter that I call "max_divergence"; see the section "**2 Which parameters to optimize?**".
3. While the optimization of "-s" by Mabs-hifiasm is performed directly, Mabs-flye log-transforms "max_divergence" and performs the golden section optimization for $\log_{10}$("max_divergence") due to the nature of "max_divergence". Basically, if a user provides Mabs-flye with very accurate reads (for example, with an error rate of approximately 1%), then fine-tuning of "max_divergence" may be beneficial. On the other hand, if a user provides Mabs-flye with highly inaccurate reads (for example, with an error rate of approximately 15%), then the parameter tuning should be more "coarse-grained". This is achieved by logarithmically transforming "max_divergence". While the interval of "-s" is [0; 1], the interval of "max_divergence" examined by Mabs is [0.0001; 0.5] or, in other words, [0.01%; 50%].
4. When optimizing "-s", Mabs-hifiasm assembles the whole genome, but Mabs-flye assembles only genes.
   Hifiasm is a fast assembler, especially taking into account that it can reuse intermediate files to produce an assembly with another "-s". On the other hand, Flye needs to perform an assembly for each "max_divergence" from the very beginning, which makes it slow. To address this problem of Flye, Mabs-flye uses only "BUSCO reads" (for the definition, see above) during assembly. Thus, Mabs-flye assembles only genes and evaluates AG only for genes. When the optimal "max_divergence" is found, Mabs-flye performs the final assembly, this time using all reads.

Assembling only genes may have a potential downside since BUSCO reads are reads that align to ancestral BUSCO proteins; thus, if the genome being assembled has very long introns and reads used for assembly are relatively short, introns may not be fully covered. This will lead to fragmentation of genes in the assembly of Mabs-flye, which in turn leads to problems in properly calculating the number of BUSCO genes and, thus, detrimentally affects the calculation of AG. However, typical genomic Oxford Nanopore reads in 2023 have lengths on the order of 10 kbp, which is probably more than the typical length of eukaryotic introns (JiaYan *et al.*, 2013). Thus, assembling only genes to find the optimal "max_divergence" is probably rational.

5. In contrast to Mabs-hifiasm, Mabs-flye polishes genes using Proovframe.
   A problem with error-prone reads is that an assembly made from them will also have many errors. Such errors are usually insertions or deletions of several bases (Kundu *et al.*, 2019; Huang *et al.*, 2021; Wick *et al.*, 2021). As they occur in CDSs, they likely lead to frameshifts, thus harming the ability of Mabs to find BUSCO genes and, thus, to calculate AG.
   One way to deal with this problem is to polish the assembly with accurate short reads. Alignment-based polishers, such as Racon (Vaser *et al.*, 2017), Pilon (Walker *et al.*, 2014) and POLCA (Zimin and Salzberg, 2020), will considerably increase the computational time of Mabs-flye since polishing is required for each of the 10 points tested by the golden section method, and read alignment is a time-consuming operation. An alternative is to use alignment-free polishers, such as ntEdit (Warren *et al.*, 2019), which are faster. However, alignment-free polishing is not as accurate as alignment-based polishing.
   A simple alternative is to use pseudopolishing by Proovframe (Hackl *et al.*, 2021). Proovframe is a tool that aligns reference proteins to a genome assembly and fixes places that cause frameshifts. This allows us to fix frameshifting assembly errors, thus making the detection of BUSCO genes possible. Mabs uses ancestral BUSCO proteins as the reference for Proovframe.
   Polishing by Proovframe has a downside in that the actual genome sequence may be distorted since it is fixed based on sequences of ancestral proteins that likely differ from sequences of current proteins of this species. For example, a frameshift in an actual pseudogene in a studied genome will be removed by Proovframe. For this reason, when the optimal "max_divergence" has been determined and Mabs-flye makes the *final* assembly using *all* reads, Proovframe is not used.

### 4 Avoiding biases during the development and testing of Mabs

#### 4.1 General considerations

A number of biases may lead to exaggeration of the quality of a bioinformatic program. Here, I describe how I dealt with two relatively nonobvious biases.

#### 4.2 Avoiding overfitting of Mabs to specific genomes

If, during its development, a genome assembler is tested on some genomes, the algorithm of this assembler may become overfitted to produce good results for these particular genomes and these particular reads. If the assembler is then compared with other assemblers on the same genomes and reads, it may outperform them, but this will not mean that the studied assembler will outperform them for other genomes and reads.

To address this problem, during the development of Mabs-hifiasm and Mabs-flye, I tested their ability to assemble genomes other than those used for comparison with Hifiasm and Flye in this article. Namely, during development, I used genomes of *Arabidopsis thaliana* (PacBio HiFi and

Oxford Nanopore reads), *Caenorhabditis elegans* (PacBio CLR reads) and the *Fagopyrum esculentum* cultivar Dasha (PacBio HiFi and Oxford Nanopore reads).

The first two genomes are small (approximately 100 Mbp) and allow for quick testing of Mabs, although their assemblies usually had no haplotypic duplications at all. On the other hand, the genome of *Fagopyrum esculentum* represents an ideal case to test a genome assembler. During the last million years, *Fagopyrum esculentum* experienced a fast expansion of 10 kbp-long transposable elements that tripled its genome, increasing the genome size from approximately 500 Mbp to approximately 1.5 Gbp (Penin *et al.*, 2021). Considering the recentness of this transposable element explosion, their copies are similar to each other, thus increasing the difficulty of genome assembly. Additionally, samples of the cultivar Dasha that were used for production of PacBio HiFi and Oxford Nanopore reads had relatively high heterozygosity (approximately 4%, to be published), which also increases the complexity of the assembly. Thus, the *Fagopyrum esculentum* cultivar Dasha represents a difficult case for genome assembly and, consequently, is highly suitable for tuning genome assemblers. The creation of a high-quality assembly of *Fagopyrum esculentum* is underway.

##### 4.3 Avoiding circular reasoning

Mabs uses the detection of BUSCO genes during assembly. The quality of assembly of BUSCO genes (i.e., "AG") is maximized by both Mabs-hifiasm and Mabs-flye. On the other hand, in Fig. 1B and Supplementary Table S3, AG is reported as the metric of assembly quality. Additionally, in Supplementary Table S3, BUSCO results are reported for the same purpose. This may create a bias in that the same metric is used to assess the assembly as is maximized during the assembly. To address this, I split all BUSCO datasets into two parts: the first part was used by Mabs-hifiasm or Mabs-flye to calculate AG during the assembly, while the second part was used to calculate AG for Fig. 1B and Supplementary Table S3 and perform BUSCO analyses for Supplementary Table S3 (see Supplementary Table S5).

#### 5. Used programs and their parameters

##### 5.1 Preprocessing of reads

To accelerate the assembly, long reads that provided genome coverage above 50 were downsampled to coverage 50 by Filtlong 0.2.1 (Wick, 2017). Quality trimming and adapter trimming of Illumina reads were performed by Fastp 0.21.0 (Chen *et al.*, 2018) with the following criteria:

1. Adapters were trimmed using the default method of Fastp, which does not require knowledge of adapter sequences.
2. Bases with Phred quality scores below 3 were removed from the 3'-ends.
3. If a 5 bp window in a read had an average Phred score below 15, this window and everything towards the 3'-end of the read were removed.
4. If the average Phred quality score of a read remained below 20 after the abovementioned procedures, the read and its pair were removed.
5. If after the abovementioned procedures the length of a read became less than 30 bp, the read and its pair were removed.

##### 5.2 Genome assembly

Genomes were assembled with Mabs 2.11, Hifiasm 0.16.1, and Flye 2.9.1.

Hifiasm was run with default parameters. All assemblies performed by Hifiasm were made with PacBio HiFi reads, except the assembly of the genome of *Heracleum sosnowskyi*, where Hi-C reads were also used. Hifiasm has the capability of using Hi-C reads for assembly graph simplification (Cheng *et al.*, 2022).

During assembly with Flye, PacBio CLR reads of *Myripristis murdjan* were provided with the option "--pacbio-raw", while Oxford Nanopore reads of *Adineta vaga* and *Mytilus coruscus* were provided with the option "nano-raw". The option "--no-alt-contigs", which is a special option of Flye for removal of haplotypic duplication, was always used. Other parameters of Flye were default.

For Mabs-hifiasm and Mabs-flye, paths to BUSCO datasets specified in Supplementary Table S5 were provided via the option "--local_busco_dataset". By default, Mabs uses 1000 orthogroups during the assembly. The database metazoa_odb10, used for *Adineta vaga*, contains only 954 orthogroups, and a portion of them had to be used for assessing the assembly quality (see "**4.3 Avoiding circular reasoning**"). Hence, for *Adineta vaga,* the value of the parameter "--number_of_busco_orthogroups" was set to 500 instead of the default 1000. Similar to the Hifiasm assembly of *Heracleum sosnowskyi*, in the Mabs-hifiasm assembly of *Heracleum sosnowskyi,* Hi-C reads were used along with HiFi reads.

##### 5.3 Postprocessing of assemblies

For Flye assemblies, polishing was performed by HyPo 1.0.3 (Kundu *et al.*, 2019), providing coverage values of Illumina reads (the option "--coverage-short") as indicated in Supplementary Table 2. Assemblies made by Mabs-flye were polished in the same way. Hifiasm assemblies do not require polishing (Cheng *et al.*, 2021), and consequently, assemblies of Mabs-hifiasm do not require polishing either.

Deduplication was performed by Purge_dups with default parameters.

##### 5.4 Quality control of assemblies

BUSCO analysis of assemblies was performed using BUSCO 5.3.2. Datasets for the BUSCO analysis were manually constructed by excluding the orthogroups that were used by Mabs during assembly (see "**4.3 Avoiding circular reasoning**") from the datasets described in Supplementary Table S5.

The AG values of the assemblies were calculated by calculate_AG from Mabs 2.11 using the same datasets as those used by BUSCO.
